# Nanocellulose hydrogels as biointerface analogs for studying nanoparticle transport and accumulation

**DOI:** 10.64898/2026.02.02.703274

**Authors:** Joshua Prince, Darryl Taylor, A-Andrew D. Jones

**Affiliations:** Department of Civil & Environmental Engineering, Pratt School of Engineering, Duke University, Durham, NC 27708; University Program in Materials Science & Engineering, Duke University, Durham, NC 27708; Thomas Lord Department of Mechanical Engineering & Materials Science, Pratt School of Engineering, Duke University, Durham, NC 27708

**Keywords:** Biointerface materials, nanoparticles, antibiotic recalcitrance, transport phenomena

## Abstract

Nanoparticles have been proposed as drug delivery vehicles to enhance targeting and efficiency of traditional and novel therapeutics. Previous studies have identified size, surface charge, and volume exclusion as factors that influence nanoparticle diffusion and retention. However, there is little accepted or successful quantification of how these parameters influence nanoparticle penetration relative to biological adaptation and biological response. Part of the challenge is the response of living biological interfaces to many of these nanoparticle delivery vehicles and nanosized drugs. This study aimed to emulate key physicochemical barriers to diffusion found in living biomaterials by developing a tunable, synthetic hydrogel. Through the controlled exposure of 150 kDa and 2 MDa nanodextrans with neutral and negative surface charge, we evaluated the system’s ability to emulate three core physicochemical features often implicated in biofilm-associated transport resistance: size exclusion, charge interactions, and volume exclusion. We demonstrated a 30% statistically significant decrease in partition coefficients for 2 MDa nanodextran from 150 kDa nanodextran, confirming the ability of the nanocellulose-based microcaps to size based exclusion of hydrated biomaterial matrices. We combined our results with bioaccumulation data from the literature and found only the interaction of size and charge had a statistically significant effect (p-value < 0.05, F-statistic = 4.8). This controllable system can be coupled with *in silico* modeling to understand interfacial transport phenomena for nanomaterial-biomaterial interactions.

## 1. Introduction

Biological interfaces, like tissues, tumors, and microbial biofilms are more complex than engineered porous membranes[2] and subsequently the transport relationships are more complex. For example, the liquid-surface biointerface typically contains soft-layers comprised of fats, sugars, and proteins of various charges [3]. The interior of the biointerface exhibits heterogeneous porosity and tortuosity [4], heterogeneous mechanical compliance, heterogeneous hydrophilicity, and heterogeneous charge distribution due to the presence of cells and extracellular components [5]. These complex structures increase the difficulty of decoupling the impacts of charge, size, geometry, and chemistry on the transport of novel drugs and drug delivery vehicles through these biological interfaces. In microbial biointerfaces, like bacterial biofilms [6–9], this complexity is increased by the dynamic response that can occur at similar time scales to the total penetration time. The dynamic response from bacteria can include extracellular matrix production of polymers like alginate, nanocellulose, proteins like flagellin [10, 11], and secretion of extracellular DNA. These properties contribute to enhanced virulence [12] and treatment failure [13] in diseases such as cystic fibrosis and catheter associated urinary tract infections. Consequently, new therapeutic strategies, including the use of nanoparticles as drug delivery vehicles, have been proposed to enhance penetration and efficacy within biofilms[14–16].

Previous studies on the impact of mechanical stress on biointerfaces have used alginate since it is an essential component of some biofilms and has facile preparation [17–20]. However, alginate has 1 – 2 orders of magnitude higher storage and loss moduli and different chemical absorption properties [19]. We have previously shown the nanocellulose preparation used here has mechanical and certain chemical properties in a similar range as bacterial biofilms [1].

Antibiotic and nanoparticle exclusion from biofilms has been proposed to be a function of the various matrix components alginate, nanocellulose, proteins; the matrix size; commensal phage; bacterial cells, particularly their charge, and the channels in the biofilm. As heterogeneity is difficult to achieve synthetically, this may require systematic study. For example, the importance of alginate on antibody binding in *P. aeruginosa* biofilms was discovered through adding alginate back into alginate deficient mutants [18]. Recent work on diffusion in alginate showed the impact of matrix cross-linking on size-exclusion [21].

This study tested the ability of a microcap synthesis method to replicate the important effects of a biofilm on species diffusion: (a) the size-effect, (b) the volume-exclusion effect, and (c) the attachment effect. These important biofilm effects were mapped on to three hypotheses. For the size effect, the size-exclusion component was tested: do larger diffusing species accumulate at lower concentrations in the mesh domain? For the volume-exclusion effect: do diffusing species accumulate at a lower concentration in a microcap with impermeable particles compared to one without? For the attachment effect: do diffusing species accumulate at a higher concentration in the microcap when attachment sites are present?

## 2. Methods

The goal of this study is to test the ability of a synthetic biomaterial to replicate the important effects of a biofilm on species diffusion. We leveraged our recently developed nanocellulose hydrogel that has a closer mechanical stiffness to natural bacterial biofilms [1]. We embedded the hydrogel with charged and non-charged microspheres to test volume-exclusion, attachment, and charge-based exclusion. We used two sizes of nanodextrans to test size exclusion and two different functional groups and one bare nanodextran to test charge effects. We used standard optical tools for examining biofilms to measure the effects of the tested parameters, Figure 1.

**Figure 1.**
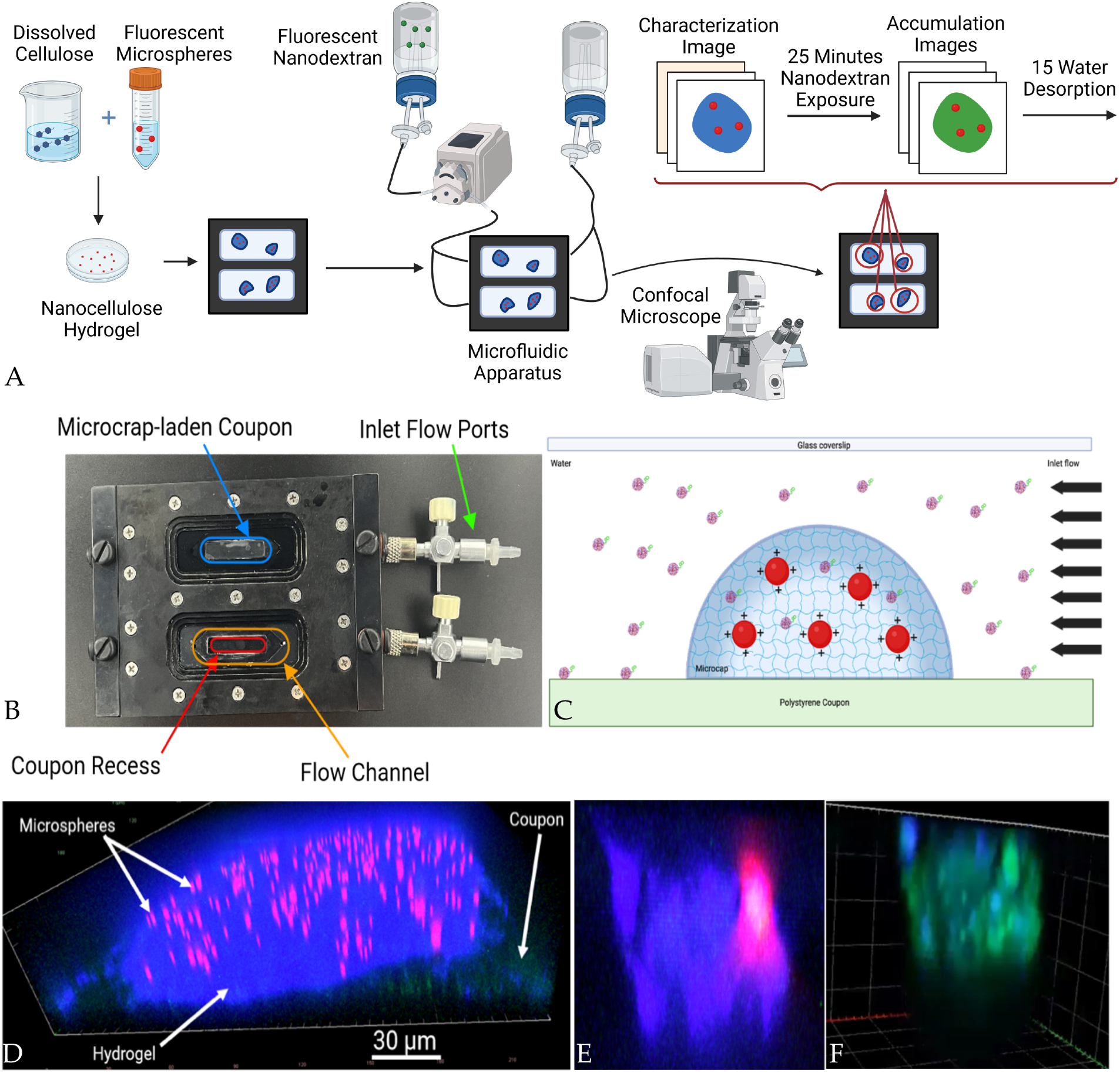
(A) Experimental protocol for testing diffusion into synthetic biofilms. Hydrogels were made as described in [1] where microspheres were added after the centrifugation step. Combined synthetic biofilms were soaked in a calcium chloride solution. (B) Resultant hydrogels were added to coupons in a flow cell (Biosurface Technology, MT, USA). (C) Schematic of flow of fluorescently-labeled nanodextran particles (green tagged, purple particles) into a matrix laden with charged fluorescently labeled polymer mircospheres (red). (C) Image of calcofluor white stained nanocellulose hydrogel (blue) with fluorescently-labeled carboxyl-coated microspheres (red) and polycarbonate coupon-microcap interface (green). (D) Image of isolated single microsphere in hydrogel before and (E) during injection with nanodextran (no modification) (green). (F)

### 2.1 Microcap Design and Nanodextran Information

To generate the microcaps, hydrogels were used as the hydrated mesh domain to achieve a size-effect. Hydrogels have been used in prior studies to simulate the extracellular matrix of the biofilm [21]. The hydrogel platform chosen for our system was a nanocellulose hydrogel. We recently developed a nanocellulose that better matched the mechanical storage and loss moduli [1].

Fluorescent polystyrene microspheres (Spherotech, IL, USA) ranging from 0.7-0.9 µm in diameter were used to simulate bacteria and achieve a volume-exclusion effect. Three distinct microspheres, each with distinct microsphere surface chemistry, were used: no surface modifications (Plain) (FP-0862-2), carboxylate-modified (Carboxyl) (FP-0862-2), and amine-modified (Amino) (FP-0862-2). The carboxylate and amine-modified microspheres were coated with these functional groups, which are either negatively or positively charged at neutral pH. These charged microsphere surfaces were designed to only act as attachment sites for the diffusing species if the diffusing species was oppositely charged. The microspheres were fluorescently-labelled with a vendor proprietary fluorescent tag.

Nanodextrans conjugated to FITC (fluorescein isothiocyanate) were used as the diffusing species for the system (TdB Labs, Uppsala, Sweden) which has been used to measure diffusion in *P. aeruginosa* biofilms [22]. Two nanodextran molecular weights used were 150 kDa and 2,000 kDa, on the premise these molecules would have different sizes, to test the size-exclusion effect. The fluorescently-labelled nanodextrans were chemically modified to have either carboxymethyl (CM) groups with negative charge (FITC-CM-dextran-150), diethylaminoethyl (DEAE) groups with positive charge (FITC-DEAE-dextran-150), or no modification at all (Dx) (FITC-Dx-150 and FITC-Dx-2000). The use of charged nanodextrans and charged microsphere surfaces was designed to produce the attachment effect.

### 2.2 Nanocellulose Solution Synthesis

A nanocellulose synthetic biofilm was generated as described in [1]. Briefly, 4.00 ± 0.05 g of dry cellulose powder (CAS#9004-34-6, Sigma-Aldrich, MA, USA) was slowly mixed in an ionic liquid solution composed of 7.00 ± 0.05 g sodium hydroxide (CAS#1310-73-2, Sigma-Aldrich, MA, USA), 12.00 ± 0.05 g urea (CAS#57-13-6, Sigma-Aldrich, MA USA), and 81.00 ± 0.05 g of reverse-osmosis filtered 18.2 MΩ water. The resulting suspension was mixed using an inert stir bar at 500 rpm until it was homogeneous and consistently cloudy. The suspension was then submerged in an ice-isopropyl alcohol bath with a temperature ranging between −10 – 0 ^O^C while still being stirred. After more than 1 hour of stirring to complete the dissolution of the cellulose, the liquid was separated into two 50 mL centrifuge tubes. The tubes were centrifuged at 10,000 rpm ( 13,751 rcf) for 10 minutes at 22 ^O^C. After centrifugation, the supernatant was decanted and saved as nanocellulose solution, with any settled solids discarded.

### 2.3 Nanocellulose-Microsphere Solutions

For each batch of nanocellulose solution made, four different nanocellulose-microsphere solutions were made: one with plain microspheres, one with carboxyl microspheres, one with amino-microspheres, and one with no microspheres. These nanocellulose-microsphere solutions were prepared by mixing 5.0±0.1 mL of nanocellulose solution with 50±0.1 µL of 1 wt% microsphere solution in the well of a 6-well culture plate. Once all four nanocellulose-microsphere solutions were prepared for each nanocellulose batch on the same plate, the plate was covered, wrapped in aluminum foil, and mixed for 24 hours on a shaking plate at 250 rpm at room temperature ranging from 22-26 °C. After 24 hours of mixing, the resulting nanocellulose-microsphere solutions were stored at 4 °C.

### 2.4 Microcap and Flow Cell Preparation

With the nanocellulose-microsphere solutions prepared, a 5 mL syringe affixed with a 27G nozzle was used for extrusion. A small amount of nanocellulose solution was extruded to prime the nozzle, the solution would then leak semi continuously. The leaking nozzle tip was pressed onto the surface of a polycarbonate coupon between 10-30 times leaving micron sized hydrogel residues. The hydrogel residues were dried at room temperature (22 – 26 °C) in a plastic petri dish for greater than 48 hours no plastic cover but aluminum foil cover, to reduce contamination and fluorescent overexposure while allowing for air flow. The resultant dried nanocellulose-microsphere composite hydrogel microcap covered coupon was loaded into a flow cell (FC 275-AL, BioSurface Technologies, Montana, USA). The flow cell chamber containing the coupon was filled with reverse-osmosis water first to rehydrate the microcap. The chamber was then filled with a 5 µM solution of Calcofluor White (CW) stain dissolved in water (29067, Biotium, California, USA). The fluorophore diffused into the microcaps, allowing for fluorescent imaging of the hydrogel domain of the microcap, Figure 1D.

### 2.5 Nanodextran Exposure and Imaging Procedure

The flow cell loaded with coupons covered with CW-stained microcaps was connected to a microfluidic apparatus with two inlets: reverse-osmosis filtered (RO) water and 30 mg/L fluorescently-labelled nanodextran dissolved in Dulbecco’s phosphate buffer solution (PBS) (Figure 1B). The flow cell was imaged on a confocal laser scanning microscope (Zeiss LSM 900, Zeiss GmBH, Oberkochen, DEU), which allowed for real-time 3D fluorescent imaging of the microcaps. Each experiment involved a three-step procedure: characterization, accumulation, and desorption. To perform an experiment, the flow chamber was flushed with 1 mL/min RO water during the characterization phase. While the flow chamber containing the microcap-covered coupon was flushed with 1 mL/min RO water (Fluigent, Massachusetts, USA), the coupons were assessed to possess at least three microcaps less than 250 µm in diameter. A z-stack series of microscope images taken at regularly spaced focal lengths to reconstruct a 3D image of a sample was taken of three microcaps on each coupon, with the three smallest microcaps selected. Field of view size was adjusted for each microcap to take up the entire view field. Nyquist sampling was used to select z-section thicknesses and image resolution. These z-stacks were set up to image three features in each microcap: the hydrogel mesh by imaging the CW stain (Laser: 405 nm/Detector Range: 410-546 nm), the impermeable microspheres (Laser: 561 nm/Detector Range: 585-700 nm), and the presence of FITC (Laser: 488 nm/Detector Range: 410-546 nm). In addition to measuring the presence of FITC, this same signal was also used to determine the location of the coupon within the image, as the coupon reflected green light at its surface. This first image before any FITC was added to the system will be referred to as the characterization image.

After the characterization image of each selected microcap, Figure 1D, the microcaps were continuously exposed to nanodextran for the accumulation phase of the experiment, Figure 1F. This was accomplished by flowing 1 mL/min of the 30 mg/L nanodextran dissolved in PBS into the flow chamber continuously for 24 minutes. After this continuous exposure to a constant concentration of nanodextran, the flow in the chamber was stopped, and the microcaps re-imaged using the same parameters as those for the characterization image, producing an accumulation image for each microcap. This was done to quantify two things: the concentration of nanodextran (via the proxy measurement of FITC-signal) in the water immediately surrounding each microcap, and the concentration of nanodextran within each microcap.

During the accumulation phase of each experiment, time-lapse imaging of a single-z-plane within one of the imaged microcaps was taken to monitor if the system reached equilibrium during each phase (which all were confirmed to reach).

### 2.6 Experimental Design

To test the functional properties of these microcaps, three predicted experimental effects were tested to determine the capacity of these microcaps to replicate the biofilm features of interest, Figure 2. The size-exclusion effect was tested to determine if a hydrated mesh was formed. The volume-exclusion effect was tested to determine if the embedded impermeable microspheres were limiting the diffusion of the nanodextran. The attachment effect was tested to determine if the diffusing species were immobilized via opposite charge interactions with the surface of the microspheres, indicating diffusing species attachment sites.

**Figure 2:**
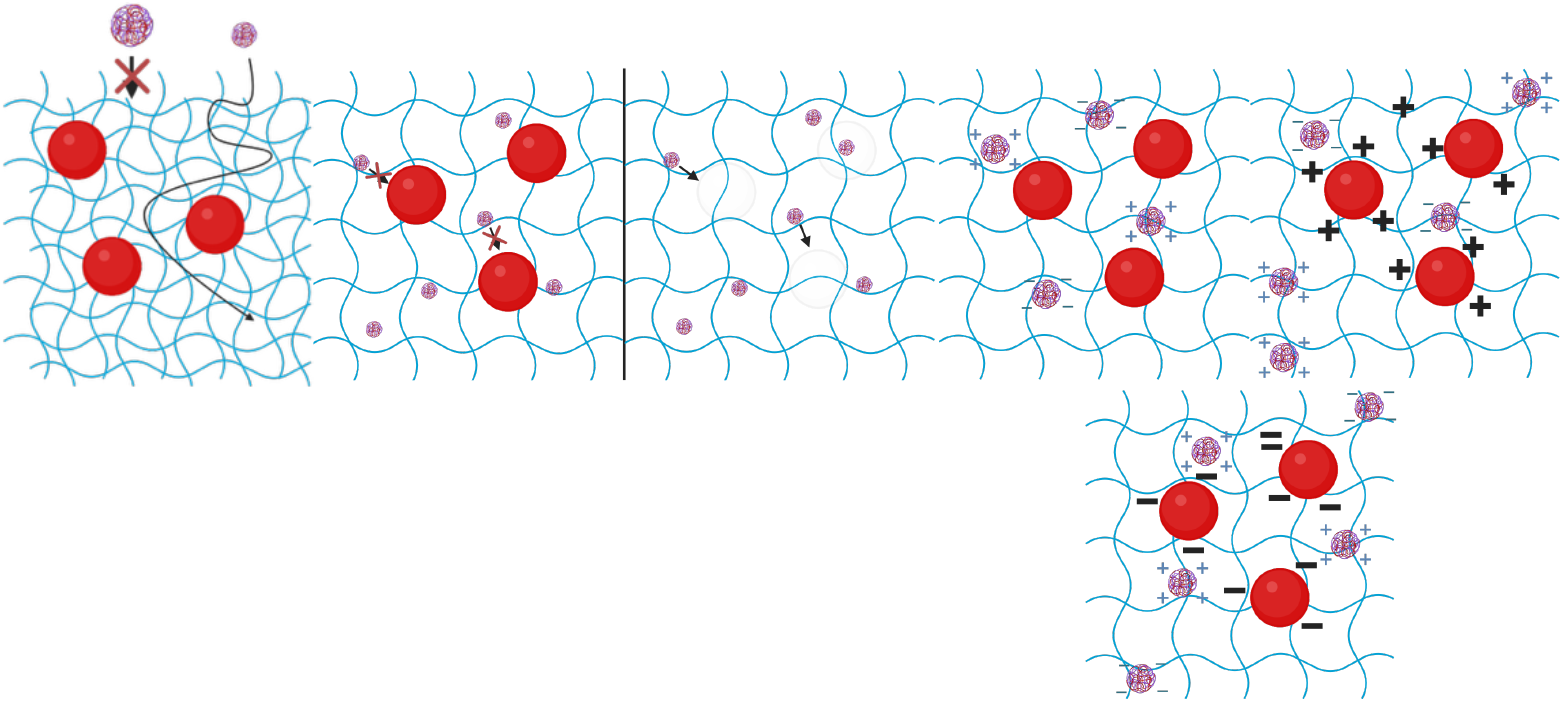
Diagrams of hypothesized (a) size-exclusion effect, (b) charge based attachment effects, and (c) volume exclusion effect.

To test for the size-exclusion effect, the two different sized nanodextrans were used: 150 kDa and 2,000 kDa. The 2,000 kDa nanodextran was expected to have a larger particle size than the 150 kDa case. This was tested and confirmed using Nanoparticle Tracking Analysis (NanoSight Pro with NTA Technology, Malvern Panalytical, Malvern, GBR) and Dynamic Light-Scattering measurements (Particle Analyzer Litesizer 500, Anton-Paar, Graz, AUT), shown in Table 1. Additional details on nanoparticle characterization are provided in the Supplementary Information. While the size and charge in Table 1 are larger and less in range, this is likely due to testing in PBS. The vendor reports the 2 MDa and 150 kDa nanodextran particles have 27 nm and 8.5 nm hydrodynamic radius respectively. Sankaran et al., reported the zetapotential same particles from the same vendor measured in deionized water 34.3±3.8 mV, −23.5±3.2 mV, −5±3 mV for diethylaminoethyl (DEAE), carboxymethyl (CM), and no modification respectively.

**Table 1:**
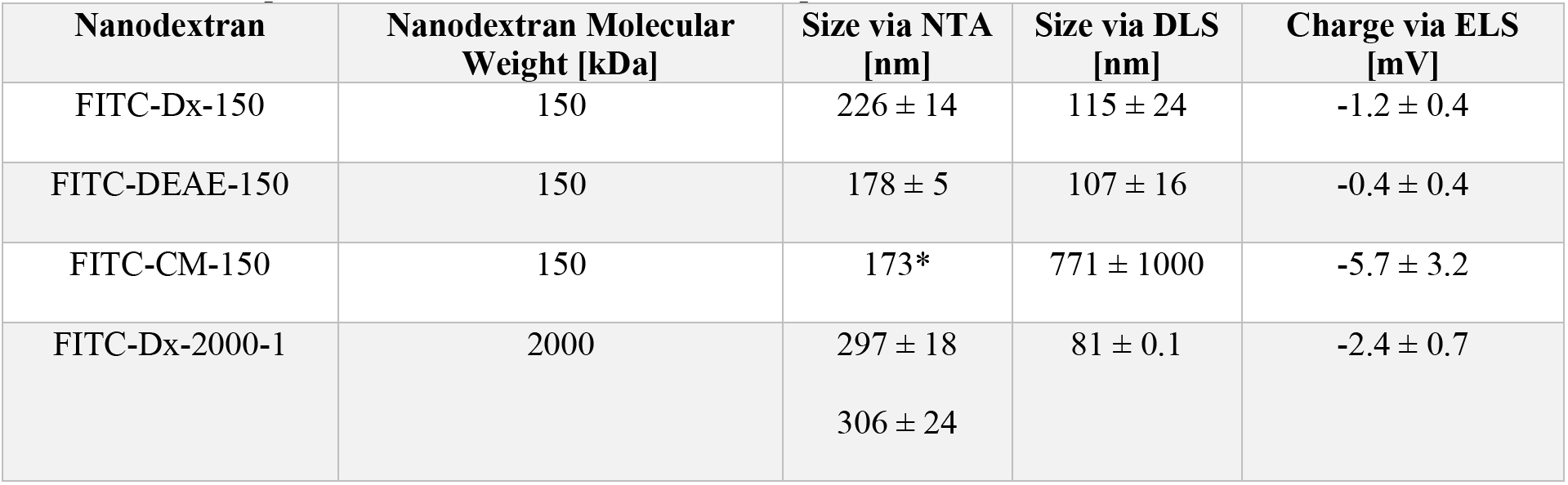
Nanodextran Characterization. All nanodextrans dispersed in Dulbecco’s Phosphate Buffer Solution (PBS). NTA error reported as standard error from measured particle size distribution (n=5). <u>*</u>-Standard error not reported on instrument. FITC-Dx-2000 size measured twice on NTA, both sizes reported. DLS/ELS error reported as standard deviation of three triplicate measurements.

| Nanodextran | Nanodextran Molecular Weight [kDa] | Size via NTA [nm] | Size via DLS [nm] | Charge via ELS [mV] |
|---|---|---|---|---|
| FITC-Dx-150 | 150 | 226 ± 14 | 115 ± 24 | -1.2 ± 0.4 |
| FITC-DEAE-150 | 150 | 178 ± 5 | 107 ± 16 | -0.4 ± 0.4 |
| FITC-CM-150 | 150 | 173* | 771 ± 1000 | -5.7 ± 3.2 |
| FITC-Dx-2000-1 | 2000 | 297 ± 18<br>306 ± 24 | 81 ± 0.1 | -2.4 ± 0.7 |

Since hydrated meshes such as a biofilm extracellular matrix and a hydrogel are comprised of disorganized, overlapping biopolymer chains, they can allow particles to diffuse through them, but only up to a certain size. As the particle reaches a characteristic “mesh size”, less of the volume within the hydrated mesh network is available for it to occupy. Once the particle reaches a critical size, no amount of particle can accumulate within the matrix. Thus, as particle size increases, the equilibrium concentration it reaches within a hydrated matrix decreases, as less volume is available for the particle to permeate into. Hence, a decrease in diffusing species concentration with particle size within the matrix at equilibrium shows the presence of a size-exclusion effect in the microcap, which in our system is likely indicative of the formation of a hydrogel. Microcaps prepared using identical conditions were used in size-exclusion experiments.

To test for the volume-exclusion effect, hydrogels were prepared either with microspheres within them or without microspheres. The microcaps without microspheres within them would be expected to reach a higher concentration of nanodextran if the volume-exclusion effect was observed. The reasoning for this is similar to the reasoning for the size-exclusion effect: certain portions of the microcap volume, specifically the volume occupied by the microspheres, would be unavailable for the nanodextran to occupy, leading to a lower concentration overall in the microcap. Identical nanodextrans were used in volume-exclusion experiments.

To test for the attachment effect, hydrogels embedded with three different microspheres were prepared and tested against three different nanodextrans. The different microspheres tested were no surface functionalization (Plain, neutrally charged surface), amine-functionalization (Amino, positively charged surface), and carboxyl-functionalized (Carboxyl, negatively charged surface). The different nanodextrans tested were not chemically modified (None, neutrally charged), carboxymethyl-modified (CM, negatively charged) and diethylaminoethyl-modified (DEAE, positively charged). Since we only expect oppositely charged combinations to lead to attachment between microsphere surface and nanodextran, we expect to see significant increases in nanodextran concentration within the microsphere only for the combination of CM-modified nanodextran accumulating in microcaps with amine-modified microspheres, and DEAE-modified nanodextran accumulating in microcaps with carboxyl-modified microspheres. Two different control cases are considered for each of these combinations: (a) charged nanodextran and uncharged microspheres and (b) uncharged nanodextran and charged microspheres. Decreases in nanodextran concentrations within the microcap would be expected in both cases compared to the oppositely charged nanodextran/microspheres case.

### 2.7 Image Analysis

To determine nanodextran concentration for each z-stack image taken, each pixel was assigned to be in one of four spatial domains in the image: (1) the water domain (liquid domain Ω*_L_*), (2) the hydrogel domain (interstitial domain Ω*_I_*) (3) the microsphere domain (Ω_0_), and (4) the coupon domain (solid domain Ω*_S_*) [23].

This was done by segmenting the hydrogel using the CW signal, the microspheres using the AF594 signal, and the coupon using the FITC signal in each image, and assuming all remaining pixels were within the water domain. The total microcap domain was the union of the microcap and microsphere domains.

The microspheres were segmented using an Otsu threshold on the AF594 signal in MATLAB. The hydrogel domain was segmented in ImageJ using an Otsu threshold, followed by a region-filling algorithm (calcofluor white stain only penetrated ∼10 µm into hydrogel), followed by an algorithm for discarding very small filled regions. The coupon was segmented using an edge-detection algorithm along the z-direction. The coupon segmentation algorithm was designed to favor a pixel as coupon as opposed to water or hydrogel, to preserve the accuracy of the water and hydrogel signal over the coupon signal.

With each pixel assigned to a domain, the average value of the nanodextran/FITC signal in each domain in the microcap was quantified. Rather than use average FITC signal in the microcap, [*FITC*]*_microcap_*, directly, the ratio of the FITC signal in the microcap to the FITC signal in the water, [*FITC*]_*water*_, or the nanodextran-microcap partition coefficient, *K_P_*, was calculated for each microcap’s accumulation image, Equation 1. This ensured any variation in FITC concentration and signal within the flow cell was normalized between microcaps.

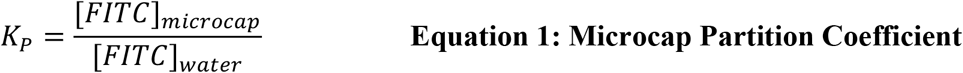

The MATLAB code for image analysis is included as Supplementary Information and is available via Github.

### 2.8 Statistical Analysis

**Figure 1:**
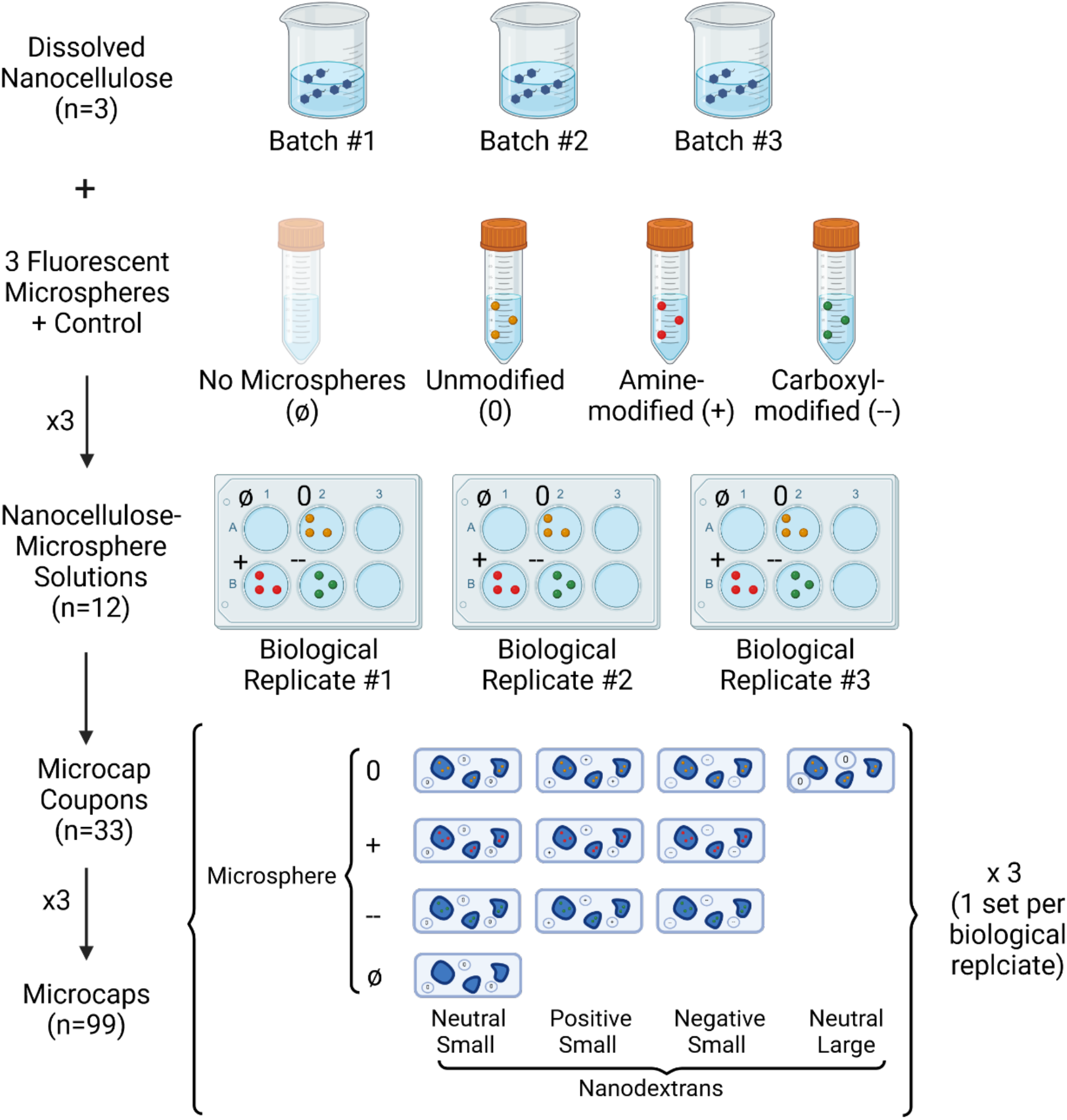
Design of Experiments and Replicates.

For each nanodextran-hydrogel combination tested, two types of replicates were used, all on the hydrogel side: batch replicates and microcap replicates. Three batches of each hydrogel tested (no microspheres, plain microspheres, carboxyl-microspheres and amine-microspheres) were generated, leading to 12 different hydrogels with three batch replicates each. For each batch, one coupon covered with microcaps was generated for each nanodextran tested, leading to 33 different coupons, with three microcap replicates on each coupon for a total of 99 microcaps tested.

To compile the data for each experimental condition (nanodextran-microcap combination), the nanodextran-microcap partition coefficient for each microcap tested was first found (see Image Analysis for how this was calculated). This value was then averaged for the three microcaps on each coupon tested, as these replicates were determined to be technical replicates, which warrant averaging together for the purposes of statistical analysis. This left three nanodextran-microcap partition coefficients for each experimental condition, each one representing a different hydrogel batch. These three nanodextran-microcap partition coefficients comprised the sample for each experimental condition.

For each set of experimental conditions compared for each hypothesis, the three nanodextran-microcap partition coefficients for each batch were compared to each other using Welch’s one-way t-tests using an alpha of 0.05. Welch’s test was used since variances were not assumed to be equal. One-way tests were used since only an effect in one direction would constitute evidence for each hypothesis. All samples passed the Shapiro-Wilks test for normality.

We tested our values and values from the literature to determine if the size, charge, size and charge, or a two-way interaction between size and charge explain bioconcentration factor using multiple regression with ANOVA. We tested three linear and one non-linear model for size, charge, size and charge, and coupled size-charge. While diffusion coefficients are a common end-point of antibiotic and nanoparticle studies [24], antibiotic effectiveness and environmental risk assessment [25, 26] are less dependent on the rate than on the total accumulation. Our non-comprehensive search identified nine independent studies had size, charge and bioconcentration factors, or the data to produce the bioconcentration factor. Two studies had size but not charge and the bioconcentration factor, and two studies whose ranges for charge or size required assuming the midpoint (see SI Table 1 for included studies and values). These studies produced 38 observations for size and 28 observations for charge and size.

## 3. Results

The microcaps generated using this synthesis procedure were tested for their ability to replicate the core features of a synthetic biofilm. These included size-exclusion effects, volume-exclusion effects and attachment effects. Different combinations of microcaps and diffusing nanodextran were used to test for these effects through equilibrium nanodextran partition coefficients in the microcaps.

### Testing for Hydrated Mesh Property via the Size Effect

To test for the size-exclusion effect, nanodextran-microcap partition coefficients for microcaps with plain microspheres were compared between a 150 kDa nanodextran and a 2,000 kDa nanodextran. The results show a statistically significant size-exclusion effect in the hydrogel mesh was observed, Figure 3A.

**Figure 3.**
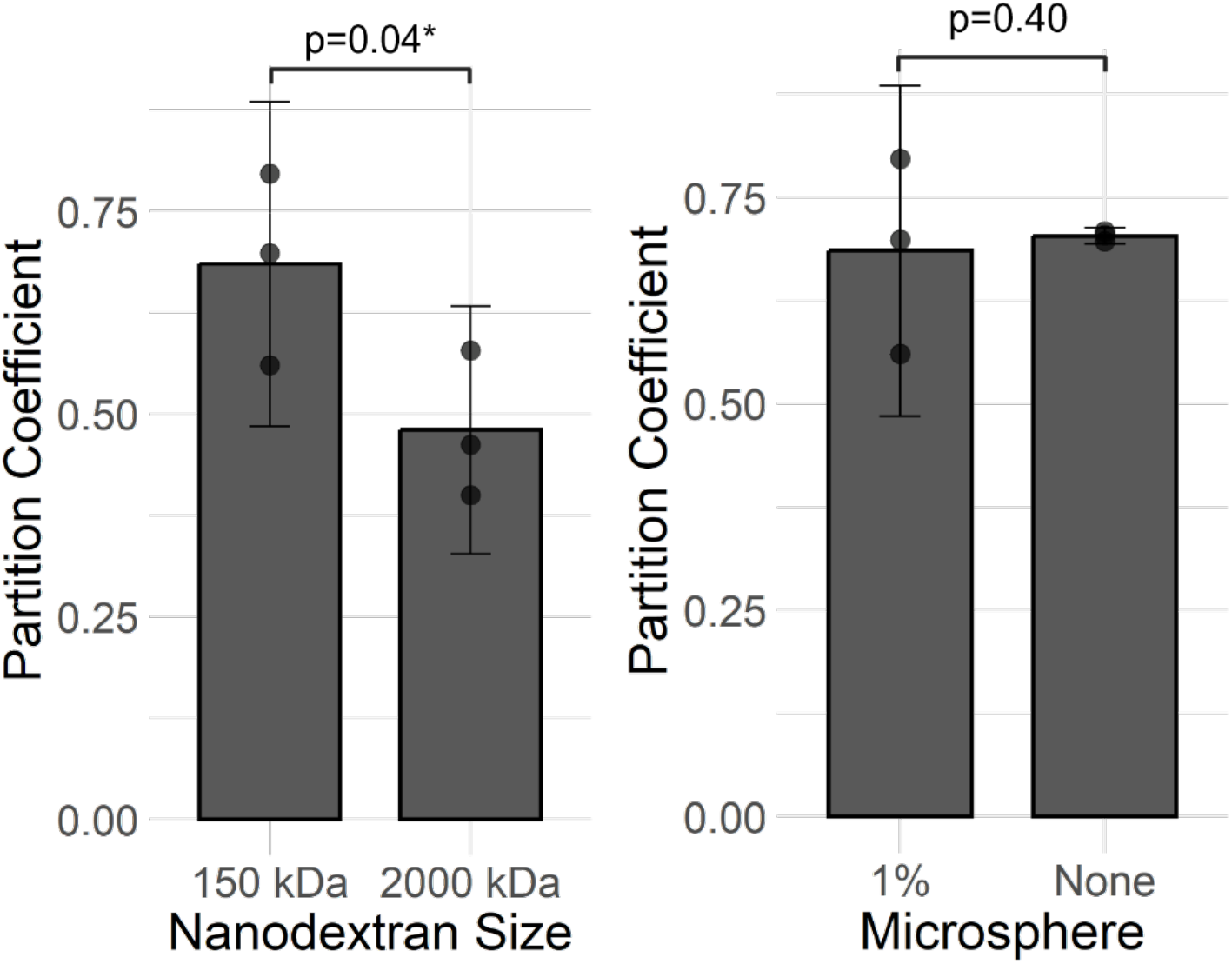
(A)Size-Exclusion Effect Test for Verifying the Hydrogel Mesh Property of the Synthetic Biofilm System. Experiments used 4 wt% hydrogels loaded with 0.01 wt% plain polystyrene microspheres with neutrally charged nanodextrans. (B) Volume-Exclusion Effect Test for Verifying the Impermeable Subdomain Property of the Synthetic Biofilm System. Experiments used 4 wt% hydrogels loaded with variable wt% plain polystyrene microspheres and neutrally charged, 150 kDa nanodextran. Partition coefficient represents nanodextran-microcap partition coefficient. Error bars represent 95% confidence intervals on the sample. P-value represents results from one-tailed Welch’s t-test.

### Testing for Impermeable Sub-volume via the Volume-Exclusion Effect

To test for the volume-exclusion effect, nanodextran-microcap partition coefficient for 4 wt% hydrogel microcaps embedded with either no microspheres or 0.01 wt% plain microspheres were compared using a neutrally charged, 150 kDa nanodextran. The results, as shown in Figure 3B, show no volume-exclusion effect due to the presence of the microspheres was observed.

### Testing for Attachment Sites via the Attachment Effect

To test for the attachment effect, 4 wt% hydrogels embedded with 0.01 wt% neutrally, negatively, and positively charged microspheres were tested against neutrally, negatively, and positively charged 150 kDa nanodextrans, Figure 6. Opposite charge combinations were tested for increases in nanodextran-microcap partition coefficients compared to both the same charged nanodextran and uncharged microspheres and the uncharged nanodextran and same charged microspheres. No statistically significant attachment effect was observed across all comparisons. A non-statistically significant attachment effect was observed for the microcaps embedded with amine-modified microspheres with a carboxymethyl-modified nanodextran for comparisons to both control cases.

**Figure 4:**
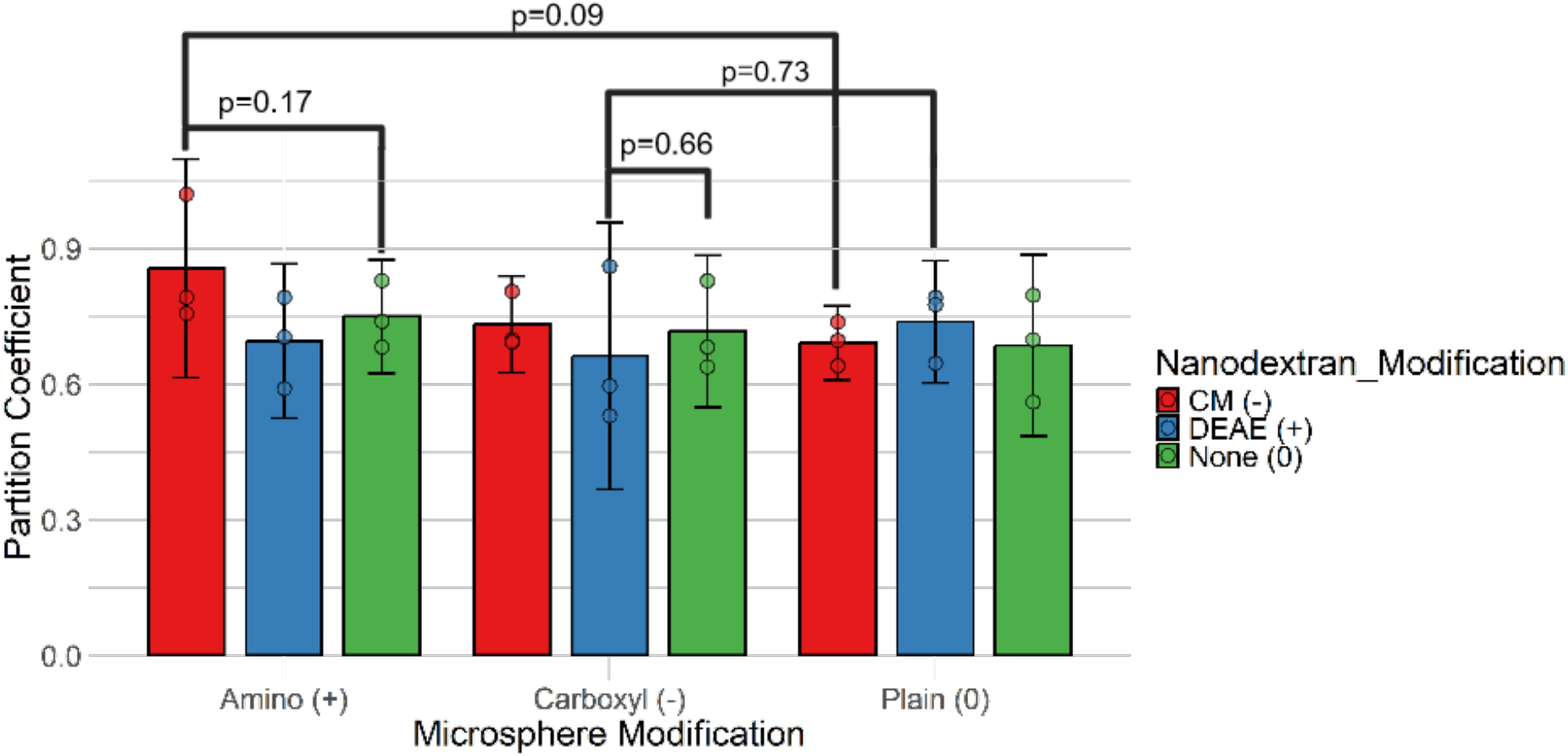
Attachment Effect Test for Verifying Presence of Attachment Sites in the Synthetic Biofilm System. Experiments used 4 wt% hydrogels loaded with variable 0.01 wt% plain-, carboxyl- and amine-modified polystyrene microspheres and neutrally, positively or negatively charged, 150 kDa nanodextrans. Partition coefficient represents nanodextran-microcap partition coefficient. Error bars represent 95% confidence intervals on the sample. P-values represent results from one-tailed Welch’s t-test.

We tested our values and values from the literature to determine if the size, charge, or both could explain the variance in bioconcentration factor using multiple regression, Table 2.

**Table 2.** ANOVA results of this study compared with literature values.

| **Analysis** | **Model** | **N** | **R²** | **F-statistic** | **p-value** |
|---|---|---|---|---|---|
| **Size alone\*** | pBCF = *β*<sub>0</sub> + *β*<sub>1</sub>*x*<sup>\*</sup> + *ε* | 38 | 0.024 | F(1,36) = 0.895 | 0.351 |
| **Charge alone** | pBCF = *β*<sub>0</sub> + *β*<sub>1</sub>*z*<sup>\*</sup> + *ε* | 28 | 0.025 | F(1,26) = 0.661 | 0.423 |
| **Size and Charge** | pBCF = *β*<sub>0</sub> + *β*<sub>1</sub>*x*<sup>\*</sup> + *β*<sub>2</sub>*z*<sup>\*</sup> + *ε* | 28 | 0.151 | F(2,25) = 2.229 | 0.129 |
| **Size–charge interaction** | pBCF = *β*<sub>0</sub> + *β*<sub>1</sub>*x*<sup>\*</sup> + *β*<sub>2</sub>*z*<sup>\*</sup> + *β*<sub>3</sub>*x*<sup>\*</sup>*z*<sup>\*</sup> + *ε* | 28 | 0.376 | F(3,24) = 4.816 | 0.00917 |

**N number of independent observations, *R*<sup>2</sup>fitting parameter. \*Includes two studies (Flood & Ashbolt[27] and Miller et al [28]) that did not report charge. Both parameters are the centered log,*x*<sup>\*</sup> = *x* − *x̄*, and *z*<sup>\*</sup> = *z* − *z̄*, where *x* = log<sub>10</sub>(*D*) with the diameter *D* in nm and *z* = log<sub>10</sub>(*ζ* + 61) with the *ζ*–potential in mV.**

While size is generally reported to be important for transport into biofilms, we found no significant effect in the range of size from 22 – 1500 nm. While some studies have reported significant effects for charge even in mixed community biofilms, consistent with what was measured in this study, we found no significant effect in the range of −60 to +34.3 mV. This analysis does support a combination of size and charge effects bioconcentration of nanomaterials.

## 4. Discussion

This research sought to create a synthetic biofilm system for use studying species transport in biofilms. To accomplish this, nanocellulose hydrogel microcaps loaded with polystyrene microspheres were developed to proxy bacterial biofilms in the important features of a hydrated mesh, an impermeable subdomain, and presence of attachment sites. The replication of these features in the generated microcaps was tested for by quantifying an important experimental effect for each feature: the size-exclusion effect for verifying the hydrogel mesh, the volume-exclusion effect for verifying the impermeable subdomain, the attachment effect for verifying presence of attachment sites.

Experimental results indicated that the size-exclusion effect was verified, and therefore the hydrogel mesh feature of the microcaps. However, the attachment effect and the volume exclusion effect were not observed. This indicates that the microcaps as currently designed are unable to replicate the key biofilm properties needed to perform transport studies of an impermeable subdomain and presence of attachment sites.

This finding of a size-effect agrees with prior literature on chemical species diffusion in single bacterial species biofilms [29, 30]. However, evidence of this effect is limited in multispecies biofilm literature on nanoparticles [31]. This is most likely due to effects of agglomeration, dissolution and changes to the nanoparticle corona in multispecies biofilm studies having strong effects on effective nanoparticle size during diffusion [32, 33]. The lack of a clear charge effect is common in the literature on diffusion of chemical species in hydrogel matrices such as biofilms [22, 23, 34–36]. While not confirmed by our experiments, this is likely confirmed by our multiple regression analysis of values in the literature showing that the interaction between size and charge was a stronger predictor of bioconcentration than size and charge alone. While this result is anticipated due to the aforementioned and well known corona, the focus in the literature of modifying one assuming independence of the other has likely limited the expected results. Furthermore, we have previously reported a coupled interaction between size and charge for antibiotic interaction with biofilm binding sites and biofilm porosity.[5].

### 4.1 Limitations

A null effect for volume-exclusion and attachment is potentially a limitation of the study platform. The use of a 0.01 wt% microsphere concentration within the hydrogels was possibly too low to observe the anticipated effects. If little volume is made impermeable due to low concentration of microspheres, then the volume-exclusion effect would be anticipated to be below a statistically significant threshold. Furthermore, too low of a concentration of microspheres would mean there are fewer sites for nanodextran attachment, also producing an attachment site effect below any statistically significant threshold. As Tseng et al. showed [37] and we previously confirmed [5], biofilms produce specific binding sites for specific drugs independent of charge, so while we tested charge-charge attachment effects, this is a poor descriptor of all possible attachment effects.

Assuming the hypothesis that the non-observed effects were primarily due to a low concentration of microspheres within the microcaps, further experimental work could replicate these experiments with higher concentrations of microspheres and look for the same effects on attachment and volume-exclusion. Further evidence for the size-exclusion effect could be found by comparing current results to higher weight percentage nanocellulose hydrogels, which theoretically lead to smaller critical mesh size, which would be anticipated to cause lower nanodextran concentrations in these higher weight percentage microcaps [29, 38].

## 5. Conclusion

This study aimed to emulate key physicochemical barriers to diffusion found in natural biofilms by developing a tunable, synthetic microcap system. Through the controlled exposure of nanodextrans with varying size and surface charge, we evaluated the system’s ability to emulate three core physicochemical features often implicated in biofilm-associated transport resistance: size exclusion, charge interactions, and volume exclusion. The results demonstrated a statistically significant size-exclusion effect, confirming the ability of the nanocellulose-based microcaps to mimic the selective permeability of hydrated biofilm matrices. However, the designed system did not display statistically significant volume-exclusion or attachment effects, suggesting that the current microsphere concentration and charge configurations were insufficient to replicate these additional features.

These findings reflect patterns observed in the literature on natural biofilms, where size-based diffusion hindrance is commonly reported, but charge-based interaction and volume exclusion are more context-dependent. We confirmed this with a regression analysis of nine independent studies showing that only a coupled size-charge interaction explained bioconcentration.

Future studies should explore increased microsphere loading, use of more biologically relevant surface chemistries to better recapitulate these additional transport-limiting features. Ultimately, refining this synthetic biofilm platform will enhance its utility in advancing the rational design of nanoparticle-based antibiofilm therapies.

## CRediT Statement

JP: Conceptualization, Methodology, Data curation, Formal Analysis, Visualization, Investigation Writing-Original draft preparation. DT: Methodology, Resources, Writing – Reviewing and Editing. A-Andrew Jones: Conceptualization, Supervision, Writing – Reviewing and Editing Methodology, Project Administration, Funding Acquisition, Formal Analysis

## Acknowledgements

- DT was supported in part by the National Science Foundation under Grant No. DGE-2022040.
- Research reported in this publication was supported by the National Institute of General Medical Sciences of the National Institutes of Health under Award Number NIH R35 GM142898 The content is solely the responsibility of the authors and does not necessarily represent the official views of the National Institutes of Health.

