## Supplementary Tables and Methods for "Nanocellulose hydrogels as biointerface analogs for studying nanoparticle transport and accumulation"

**Supplementary Information** Nanocellulose hydrogels as bio-interface analogs for studying nanomaterial transport and accumulation

Joshua Prince[1,a], Darryl Taylor[2,b] and A-Andrew D. Jones, III[1,2,3,c,*]

**Affiliations:**

1. Department of Civil & Environmental Engineering, Pratt School of Engineering, Duke University, Durham, NC 27708
2. University Program in Materials Science & Engineering, Duke University, Durham, NC 27708,
3. Thomas Lord Department of Mechanical Engineering & Materials Science, Pratt School of Engineering, Duke University, Durham, NC 27708

**Nanoparticle Characterization**

Three methods were used to analyze the nanodextrans in the study. The first was Nanoparticle Tracking Analysis (NTA), performed using a Malvern NanoSight Pro with a 532 green laser, which detects particle size using light microscopy to track individual particle movements. The second was Dynamic Light Scattering (DLS), performed using a Anton-Paar Litsizer 500 Particle Analyzer, which detects particle size using fluctuations in light signal through a sample due to Brownian motion. The final was Electrophoretic Light Scattering (ELS), which detects particle surface charge using fluctuations in light signal through a sample due to electrophoresis.

All samples were prepared by taking dry powder formulations of the nanodextrans and dispersing them in Dulbecco's Phosphate Buffer Solution (0.137 M sodium chloride (CAS# 7647145, ACS reagent grade ≥99.0% Sigma-Aldrich), 0.0027 M potassium chloride (CAS# 7447407, ACS reagent grade 99.0 – 100.5%, Sigma-Aldrich), 0.01 M sodium phosphate dibasic (CAS# 7558794, bioreagent grade ≥99.0% Sigma-Aldrich), 0.0018 M potassium phosphate monobasic (CAS# 7778770, ≥99.0%, Sigma-Aldrich).

For DLS, 1 mL of sample was added to a plastic cuvette and analyzed. Instrument settings were a refractive index of 1.45 for the nanodextran, a refractive index of 1.33 for the PBS solution, 1 minute of equilibration time, a temperature of $25^{O}$C, back-scattering, automatic filters, automatic focus, automatic number and length of runs, and automatic analysis modules to infer diffusivity and hydrodynamic diameter distribution from generated autocorrelation functions. All samples were ran in triplicate.

For ELS, 0.600 mL of sample was added to a zeta-potential Omega cuvette (#225288) and analyzed. Instrument settings were a PBS solvent, a temperature of $25^{O}$C, 1 minute of equilibration time, automatic power adjustments, automatic number of runs, and zeta-potential inferred using the Smoluchowski relation with a Henry Factor of 1.5. All samples were run in triplicate.

For NTA, samples were analyzed using automatic equipment settings. 1 mL of sample was analyzed using a constant flowrate of 3 µL/min, with 5 replicates measurements per sample.

**Materials and Equipment**

The following chemicals were used in this experimental process:

Cellulose powder (CAS#9004-34-6 | Sigma-Aldrich | Burlington, MA);

Sodium hydroxide (CAS#1310-73-2 | Sigma-Aldrich | Burlington, MA);
Urea (CAS#57-13-6 | Sigma-Aldrich | Burlington, MA)
Sodium chloride (CAS# 7647145, ACS reagent grade ≥99.0% Sigma-Aldrich | Burlington, MA)
Potassium chloride (CAS# 7447407, ACS reagent grade 99.0 – 100.5%, Sigma-Aldrich | Burlington, MA)
Sodium phosphate dibasic (CAS# 7558794, bioreagent grade ≥99.0% Sigma-Aldrich | Burlington, MA)
Potassium phosphate monobasic (CAS# 7778770, ≥99.0%, Sigma-Aldrich)

Polystyrene microspheres (Spherotech | Lake Forest, IL) with no surface modifications (Prod# FP-0862-2),
Polystyrene microspheres carboxylate-modified (Prod# FP-0862-2)
Polystyrene microspheres amine-modified (Prod# FP-0862-2)
Fluorescent nanodextrans (TdB Labs, Uppsala, Sweden) with no chemical modification
Fluorescent nanodextrans lower molecular weight (Prod# FITC-Dx-150)
Fluorescent nanodextrans carboxymethyl groups (Prod# FITC-CM-dextran-150)
Fluorescent nanodextrans diethylaminoethyl groups (Product # FITC-DEAE-dextran-150)
Fluorescent nanodextrans with no chemical modification and high molecular weight (Prod# FITC-Dx-2000)
Calcofluor White (CW) stain (Cat# 29067 | Biotium | Fremont, CA, USA).

The following materials were used in this experimental process:
6-well culture plate (Pro# 10062-892 | VWR | Radner, PA, USA)
5 mL syringe with Leur-lock tip (SKU# 309646)
27G nozzle (SKU# NZ3270005001 | CELLLINK | Gothenburg, Sweden)
two-channel microscopy flow cell (Prod# FC 275-AL | BioSurface Technologies | Bozeman, MT, USA) with 6x25x2 mm polycarbonate coupons
1/8” ID PVC tubing (Prod# 13-712-77 | Fisher | Pittsburgh, PA).

The following equipment was used in this process:
Refrigerated centrifuge with fixed angle rotor (Prod# 5910 Ri | Eppendorf | Hamburg, Germany);
Pressure-driven microfluidic system (Fluigent | Lowell, MA) with a primary pressure controller (Prod# LU-FEZ-0345), valve controller (Prod# ELU-SEZ), computer link (Prod# LU-LNK-0002), flowrate sensor/controller (Prod# FLU-L-D), multi-switch valve (Prod# ESSMSW003) connected via 0.020” ID 1/16” OD tubing (Prod# CTQ-KIT-HQ)
Zeiss LSM 900 confocal microscope (Zeiss | Oberkochen, Germany)
NanoSight Pro with NTA Technology (Model # HBG5000 | Malvern Panalytical | Malvern, United Kingdom)
Anton-Paar Particle Analyzer Litesizer 500 (Anton-Paar | Graz, Austria)

**Image Analysis MATLAB Code**
**Also available on Github https://github.com/Dr-Jones-SEEL-Team/Bioaccumulation_NP**

Below is an example of the code used to process the z-stack imaging data in MATLAB. Separate code as outlined in the Main Body was used for ImageJ processing.

Nanodextran Accumulation in Nanocellulose Time-Lapse Imaging Data Analyzer-

**Refresh MATLAB Environment**

```
clc
close all
clear all
```

**Launch Parellel Computing Pool**

```
% parpool('local',10); %Enable parellel pool
```

**Change MATLAB path and Add bfmatlab directory to path**

```
% % Josh's Mac homepath
% homepath='/Users/joshuaprince/Library/CloudStorage/Box-Box/Quantum Biofilms/Processed Data/Synthetic-Quantum Biofilms/JP_190/JP_177_Chan1_Microcap1'; %Define home directory for code
% cd(homepath) %Change directory to this home path

% Josh's PC homepath
homepath='C:\Users\joshu\Box\Quantum Biofilms\Processed Data\Synthetic-Quantum Biofilms\JP_218\JP_218_Microcap1'; %Define home directory for code
cd(homepath) %Change directory to this home path

bfmatlabdir='bfmatlab'; %Specify directory to add
addpath(genpath(bfmatlabdir)); %Add the directory

functionscriptdir='subfunctions_v3'
addpath(genpath(functionscriptdir)); %Add directory
```

Analysis of Characterization Image

**Read in the Characterization/Accumulation/Desorption Image**

```
% file_path1='JP_190_Nanocellulose_Plain-PS_FITC-DEAE-150_20240707_Confocal_1.czi' %Load in Characterization Image
file_path2='JP_218_Nanocellulose_Plain-PS_FITC-DEAE-150_20240719_Confocal_5.czi' %Load in Accumulation Image
% file_path3='JP_177_Nanocellulose_Amino-PS_FITC-DEAE-150_20240413_Confocal_9.czi' %Load in Desorption Image
% file_paths={file_path1;file_path2;file_path3 ;
```

**Retrieve relevant metadata**

```
reader = bfGetReader(file_path2);
metadata = retrieve_metadata(file_path2);
numX=metadata(1); % image width, pixels
numY=metadata(2); % image width, pixels
numZ=metadata(3); % number of Z slices
numC=metadata(4); % number of channels
numP=metadata(5); % number of total planes
```

```
numI=metadata(6); % number of total images
locZ1=metadata(7); %Get top z-position (yes, everything is inverted in Z, might fix in post, might not, whatever)
locZ2=metadata(8); %Get bottom z-position (yes, everything is inverted in Z, might fix in post, might not, whatever)
locXc=metadata(9); %Get z-stack x-position (Center)
locYc=metadata(10); %Get z-stack y-position (Center)
sizeX=metadata(11); %Get the area covered by X-pixel
sizeY=metadata(12); %Get the area covered by X-pixel
sizeZ=metadata(13); %Get the area covered by X-pixel
wavelengthC1=metadata(14); %Get the emission wavelength for Channel 1
wavelengthC2=metadata(15); %Get the emission wavelength for Channel 2
wavelengthC3=metadata(16); %Get the emission wavelength for Channel 3

scaling= [sizeX,sizeY,sizeZ]; %Define the size of pixels in each spatial dimension
scaling=double(scaling); %Convert between MATLAB data formats
scaling_base=min(scaling);  %Define the base of the new scaling vector
for i=1:length(scaling)
   scaling(i)=round(scaling(i)/scaling_base,0);
end
zfactor=scaling(3); %Define factor needed to rescale the z-direction
%Leave room for other metadata I need to retrieve
```

**Create Master Pixel-Value Data Matrices**

```
for j=1:numC %Loops through number of channels on the file
   master_pixel_generator(file_path2,numX,numY,numZ,j); %Generate master pixel value matrix for Channel 1
end
```

So, after trying to just load in the data as is into a giant array, it is pretty clear that this will be too much of a drain on memory. Which makes sense. These are large arrays of data. So, I'll have to be more clever about only ever calling subsets of data when using it. Should be possible.

After moving over to running this script on a DCC virtual machine with boatloads of memory (but probably not enough CPUs, only 4), I was able to create these large data matrices.

**Segmenting Microspheres Channel**

Goal is to generate a binary file partitioning pixel data matrixinto "in-microsphere" volume (binary value 1) and "non in-microspeheres volume" (binary value 0).

```
datafilename=sprintf('datac3.mat');
microsphere_segmentation(datafilename)
```

**Segmentation of Channel 2: Hydrogel**

```
% Convert ImageJ  segementations into MATLAB .mat datafiles
coupon_zposition=ImageJ_convert(file_path2) %Save the z-plane with the largest amount of segmented hydrogel, use as coupon interface
```

**Segmentation of Channel 1: Coupon**

```
% Segment Coupon
```

```
datafilename=sprintf('datac1.mat');
coupon_segmenter(datafilename,coupon_zposition); %Create coupon volume below coupon
position

% Remove microsphere volume from hydrogel segmentation
datafilename_hydrogel=sprintf('datac2_ImageJsegmented.mat');

%Remove coupon volume from hydrogel segementation
datafilename_coupon=sprintf('datac1_couponsegmented.mat');
mask_remover_binary(datafilename_hydrogel,datafilename_coupon)
```

**Define water volume**

```
datafilename_coupon=sprintf('datac1_couponsegmented.mat');
datafilename_microspheres=sprintf('datac3_OTSUed.mat');
datafilename_hydrogel=sprintf('datac2_ImageJsegmented_removebinarymask.mat');
water_segmenter(datafilename_hydrogel,datafilename_microspheres,datafilename_coupon)
%Define water volume

% 3-D viewer of water segmentation

datafilename=sprintf('water.mat')
downF=2;
new_numX=round(numX/downF,0);
new_numY=round(numY/downF,0);
new_numZ=round(numZ/downF,0);
zstack_resizer(datafilename,new_numX,new_numY,new_numZ)
datafilename_resized=sprintf('water_resized.mat');
threeD_rendering(datafilename,scaling) %3-D Render the segemented volume in
volumeViewer
```

**Initialize Output Vector**

```
%Define the three entries in the hdyrogel vector
stages={'Accumulation' ;

%Define Image Phases of Interest
phases={'Microcap';'Hydrogel';'Microspheres';'Water';'Coupon' ;

%Define variable types for table
varTypes={'double','double','double','double','double' ;

%Initiaite Output Table
```

```
export_table=table('Size',[1,5],'VariableTypes',varTypes,'VariableNames',phases,'RowNames',
stages)
```

**Quantifying Average Channel 1 values in Hydrogel (with Microspheres)**

```
datafilename_pixeldata=sprintf('datac1.mat') ;
datafilename_binarymask=sprintf('datac2_ImageJsegmented_removebinarymask.mat');
%Grab hydrogel binary mask
export_table{1,'Hydrogel' =average_value(datafilename_pixeldata,datafilename_binarymask);
```

**Quantifying Average Channel 1 values in Microsphere Volume**

```
datafilename_pixeldata=sprintf('datac1.mat') ;
datafilename_binarymask=sprintf('datac3_OTSUed.mat') ;
export_table{1,'Microspheres'
=average_value(datafilename_pixeldata,datafilename_binarymask);
```

**Quantifying Average Channel 1 values in Water Volume**

```
datafilename_pixeldata=sprintf('datac1.mat') ;
datafilename_binarymask=sprintf('water.mat') ;
export_table{1,'Water' =average_value(datafilename_pixeldata,datafilename_binarymask);
```

**Quantifying Average Channel 1 values in Coupon Volume**

```
datafilename_pixeldata=sprintf('datac1.mat') ;
datafilename_binarymask=sprintf('datac1_couponsegmented.mat') ;
export_table{1,'Coupon' =average_value(datafilename_pixeldata,datafilename_binarymask);
```

**Export Nanodextran-concentration in phases matrix**

```
writetable(export_table,'JP_218_Microcap2_v3', 'WriteRowNames',true)
```

Table S1: Studies reporting charge, size, and sufficient data to compute a bioconcentration factor. Sankaran *et al* and Prince *et al* both report a partition coefficient based on fluorescent intensity.

| Study | Type | Size (nm) | Charge | Species | log(BCF) |
|---|---|---|---|---|---|
| Flood and Ashbolt 1999[1] | Latex carboxylate | 100(-) | | Wastewater treating wetland biofilm | ~2 |
| Ferry et al 2009[2] | Gold | | 6530 mV | Estruary Biofilm | 2.18 |
| Burns et al 2013[3] | Gold polystyrene sulfonate | | 50-53 mV | Estruary Biofilm | 3.46 |
| Cleveland et al 2012[4] | Citrate capped silver | 30, 115 | -18 mV | Estruary Biofilm | 1.4 – 1.5 |
| Fabrega et al 2011[5] | Citrate capped silver | 100 – 700 | -18 mV | *Pseudomonas putida* OUS 82 | 2.05 |
| Kroll et al 2016[6] | PVP capped silver | | 117-17 mV | Benthic biofilm | 4.6 |
| Wright et al 2018 [7] | Titanium dioxide | | 150-17 mV | Wastewater treating wetland biofilm | 4 - 5.6 |
| Miao et al 2015[8] | Copper oxide | 100 – 800 | -35 – +15 mV | Periphytic biofilm | 3.6 – 4.25 |

| | | | | | |
|---|---|---|---|---|---|
| Grün et al 2016[9] | Citrate capped silver | 30, 70 | -60 mV | *Aquabacterium citratiphilum* | 0.8 |
| Nevius et al 2012[10] | sulphate capped polystyrene | 22±3 | -36.01±0.59 | *Alteromonas macleodii* low salinity artificial sea water | 6.32 |
| Nevius et al 2012[10] | sulphate capped polystyrene | 22±3 | -36.01±0.59 | *Alteromonas macleodii* high salinity artificial sea water | 6.24 |
| Nevius et al 2012[10] | sulphate capped polystyrene | 22±3 | -36.01±0.59 | *Alteromonas macleodii* high NaCl | 5.53 |
| Nevius et al 2012[10] | carboxylate capped polystyrene | 24±3 | -40.24±1.74 | *Alteromonas macleodii* low salinity artificial sea water | 5.08 |
| Nevius et al 2012[10] | carboxylate capped polystyrene | 24±3 | -40.24±1.74 | *Alteromonas macleodii* high salinity artificial sea water | 5.01 |
| Nevius et al 2012[10] | Amine capped polystyrene | 190±11 | 21.89±0.61 | *Alteromonas macleodii* low salinity artificial sea water | 5.89 |
| Nevius et al 2012[10] | Amine capped polystyrene | 190±11 | 21.89±0.61 | *Alteromonas macleodii* high salinity artificial sea water | 5.79 |
| Nevius et al 2012[10] | Amine capped polystyrene | 190±11 | 21.89±0.61 | *Alteromonas macleodii* high NaCl | 4.23 |
| Sankaran et al. 2019[11] | FITC-DEAE-150 | | 2734.3±3.8 mV | *Pseudomonas aeruginosa* PAO1 | 0.51 |
| Sankaran et al. 2019[11] | FITC-CM-150 | | 27-23.5±3.2 mV | *Pseudomonas aeruginosa* PAO1 | -0.05 |
| Sankaran et al. 2019[11] | FITC-Dx-150 | | 27-5±3 mV | *Pseudomonas aeruginosa* PAO1 | -0.10 |
| Miller et al 2013[12] | L-tyrosine polyphosphate | 500-2,500 | | *Pseudomonas aeruginosa* PAO1 | 2.36 |
| Miller et al 2013[12] | L-tyrosine polyphosphate | 500-2,500 | | *Pseudomonas aeruginosa* PAO1 | 2.23 |
| Miller et al 2013[12] | L-tyrosine polyphosphate | 500-2,500 | | *Pseudomonas aeruginosa* PAO1 | 2.58 |
| Miller et al 2013[12] | L-tyrosine polyphosphate | 500-2,500 | | *Pseudomonas aeruginosa* PAO1 | 2.26 |
| Miller et al 2013[12] | L-tyrosine polyphosphate | 500-2,500 | | *Pseudomonas aeruginosa* PAO1 | 2.96 |
| Miller et al 2013[12] | L-tyrosine polyphosphate | 500-2,500 | | *Pseudomonas aeruginosa* PAO1 | 2.74 |
| Miller et al 2013[12] | L-tyrosine polyphosphate | 500-2,500 | | *Pseudomonas aeruginosa* PAO1 | 2.68 |
| Miller et al 2013[12] | L-tyrosine polyphosphate | 500-2,500 | | *Pseudomonas aeruginosa* PAO1 | 2.96 |
| Miller et al 2013[12] | L-tyrosine polyphosphate | 500-2,500 | | *Pseudomonas aeruginosa* PAO1 | 2.70 |
| This study | FITC-Dx-150 | 115 ± 24 [via DLS] | -1.2 ± 0.4 mV | Hydrogel + neutral microspheres | -0.16 |

| | | | | | |
|---|---|---|---|---|---|
| | | 226±14 [via NTA] | | | |
| This study | FITC-Dx-150 | 115 ± 24 [via DLS] 226±14 [via NTA] | -1.2 ± 0.4 mV | Hydrogel + negative charged microspheres | -0.14 |
| This study | FITC-Dx-150 | 115 ± 24 [via DLS] 226±14 [via NTA] | -1.2 ± 0.4 mV | Hydrogel + positive charged microspheres | -0.12 |
| This study | FITC-DEAE-150 | 107 ± 16 [via DLS] 178±5 [via NTA] | -0.4 ± 0.4 mV | Hydrogel + neutral microspheres | -0.13 |
| This study | FITC-DEAE-150 | 107 ± 16 [via DLS] 178±5 [via NTA] | -0.4 ± 0.4 mV | Hydrogel + negative charged microspheres | -0.18 |
| This study | FITC-DEAE-150 | 107 ± 16 [via DLS] 178±5 [via NTA] | -0.4 ± 0.4 mV | Hydrogel + positive charged microspheres | -0.16 |
| This study | FITC-CM-150 | 771 ± 1000 [via DLS]; 173* [via NTA] | -5.7 ± 3.2 mV | Hydrogel + neutral microspheres | -0.16 |
| This study | FITC-CM-150 | 771 ± 1000 [via DLS]; 173* [via NTA] | -5.7 ± 3.2 mV | Hydrogel + negative charged microspheres | -0.14 |
| This study | FITC-CM-150 | 771 ± 1000 [via DLS]; 173* [via NTA] | -5.7 ± 3.2 mV | Hydrogel + positive charged microspheres | -0.07 |
| This study | FITC-Dx-2000-1 | 81 ± 0.1 [via DLS] 297 ± 18, 306 ± 24 [via NTA] | -2.4 ± 0.7 mV | Hydrogel + Plain microspheres | -0.32 |

Table S1 : Studies Investigating Accumulation of Nanoparticles in Single Species Biofilms.

| **Study** | **Nanoparticle(s)** | | | **Bacterial Species** | **Reported Factors Affecting Nanoparticle Accumulation/diffusion** |
|---|---|---|---|---|---|
| | **Core Composition** | **Size [nm]** | **Surface charge** | | |

| | | | | | |
|---|---|---|---|---|---|
| Peulen and Wilkinson [13] | Rhodamine, Dextrans, Silver, Carboxylated Microspheres | 0.5-67.5 | Negative, Neutral, Positive | *Pseudomonas fluorescens* | Nanoparticle size, biofilm density, surface charge, |
| Guiot *et al* [14] | Dextrans, COOH-Latex and $NH_2$-Latex | 4-55 | Negative, Neutral, Positive | *Lactococcus lactis, Stenotrophonas maltophilia* | Biofilm Density, Nanoparticle Size |
| Miller *et al* [15] | L-tyrosine polyphosphate | 500-2,500 | -- | *Pseudomonas aeruginosa* PAO1 | Supernatant Flowrate |
| Fabrega *et al* [5] | Citrate-AgNP | 65 | Negative | *Pseudomonas putida* OUS 82 | Concentration of Dissolved Organic Matter |
| Morrow *et al* [16] | Quantum Dots | 23-32 | Negative | *Pseudomonas aeruginosa* PAO1-GFP | Nanoparticle size/charge |
| Nevius *et al* [17] | Polystyrene NPs | 22-190 | Negative, Positive | *Alteromonas macleodii* | Nanoparticle charge/surface chemistry |
| Tripathi *et al* [18] | Polystyrene, QDs | 10-1,000 | Negative | *Pseudomonas aeruginosa* Boston 41501 | Nanoparticle charge/surface chemistry |
| Jing *et al* [19] | Cerium Dioxide NPs | 100 | --- | *Pseudomonas fluorescens* & *Mycobacterium smegmatis* | Bacterial Species |
| Li *et al* [20] | Quantum Dots | 7.5-24 | Negative, Neutral, Positive | *Escherichia coli* DH5α | Nanoparticle charge/ hydrophobicity |
| Xiao *et al* [21] | $nC_{60}$, fullerol, Citrate-AgNP, PVP-AgNP | 27-100 | Negative | *Pseudomonas aeruginosa (*ATCC 7700) | Nanoparticle hydrophobicity, biofilm density |
| Desmau *et al* [22] | PVP-AgNP, $SiO_2$-AgNP, $SiO_2$-$NH_2$-AgNP | ~90 | Negative, Positive | *Shewanella oneidensis* MR-1 | Nanoparticle size/charge/ hydrophobicity |
| Sankaran *et al* [11] | Nanodextran | 5-20 | Negative, neutral, positive | *Pseudomonas aeruginosa* | Nanoparticle Size |
| Zhang *et al* [23] | Nanodextran, fluorophores | 1-35 | Neutral, positive, negative | *Streptococcus mutans* | Nanoaprticle Size/Charge and Biofilm pH |
| Habimama *et al*[24] | Polystyrene | 50 | Negative | *Lactococcus lactis* | Hydrophobicity |
| Hollman *et al* [25] | Fluorescent Nanosensors | ~35 | Neutral, Positive | *P. aeruginosa, Streptococcus mutans* | Nanoparticle Size |
| Rodríguez-Suárez *et al* [26] | Polystyrene | ~64 | Positive, Negative | *P. aeruginosa* | Volume-exclusion |

1 Flood, J. A. & Ashbolt, N. J. Virus-sized particles can be entrapped and concentrated one hundred fold within wetland biofilms (Reprinted from Advances in Environmental Research, vol 3, pg 403-411, 2000). *Advances in Environmental Research* **3** (1999).

2 Ferry, J. L. *et al.* Transfer of gold nanoparticles from the water column to the estuarine food web. *Nat Nanotechnol* **4**, 441–444 (2009). https://doi.org/10.1038/nnano.2009.157

3 Burns, J. M. *et al.* Surface charge controls the fate of Au nanorods in saline estuaries. *Environmental Science & Technology* **47**, 12844–12851 (2013). https://doi.org/10.1021/es402880u

4 Cleveland, D. *et al.* Pilot estuarine mesocosm study on the environmental fate of Silver nanomaterials leached from consumer products. *Science of the Total Environment* **421**, 267–272 (2012). https://doi.org/10.1016/j.scitotenv.2012.01.025

5 Fabrega, J., Zhang, R., Renshaw, J. C., Liu, W. T. & Lead, J. R. Impact of silver nanoparticles on natural marine biofilm bacteria. *Chemosphere* **85**, 961–966 (2011). https://doi.org/10.1016/j.chemosphere.2011.06.066

6 Kroll, A. *et al.* Mixed messages from benthic microbial communities exposed to nanoparticulate and ionic silver: 3D structure picks up nano-specific effects, while EPS and traditional endpoints indicate a concentration-dependent impact of silver ions. *Environ Sci Pollut Res Int* **23**, 4218–4234 (2016). https://doi.org/10.1007/s11356-015-4887-7

7 Wright, M. V. *et al.* Titanium dioxide nanoparticle exposure reduces algal biomass and alters algal assemblage composition in wastewater effluent-dominated stream mesocosms. *Science of the Total Environment* **626**, 357–365 (2018). https://doi.org/10.1016/j.scitotenv.2018.01.050

8 Miao, L. *et al.* Effects of pH and natural organic matter (NOM) on the adsorptive removal of CuO nanoparticles by periphyton. *Environ Sci Pollut Res Int* **22**, 7696–7704 (2015). https://doi.org/10.1007/s11356-014-3952-y

9 Grun, A. Y., Meier, J., Metreveli, G., Schaumann, G. E. & Manz, W. Sublethal concentrations of silver nanoparticles affect the mechanical stability of biofilms. *Environ Sci Pollut Res Int* **23**, 24277–24288 (2016). https://doi.org/10.1007/s11356-016-7691-0

10 Nevius, B. A., Chen, Y. P., Ferry, J. L. & Decho, A. W. Surface-functionalization effects on uptake of fluorescent polystyrene nanoparticles by model biofilms. *Ecotoxicology* **21**, 2205–2213 (2012). https://doi.org/10.1007/s10646-012-0975-3

11 Sankaran, J. *et al.* Single microcolony diffusion analysis in Pseudomonas aeruginosa biofilms. *NPJ Biofilms Microbiomes* **5**, 35 (2019). https://doi.org/10.1038/s41522-019-0107-4

12 Miller, J. K. *et al.* Nanoparticle Deposition onto Biofilms. *Ann. Biomed. Eng.* **41**, 53–67 (2013). https://doi.org/10.1007/s10439-012-0626-0

13 Peulen, T. O. & Wilkinson, K. J. Diffusion of nanoparticles in a biofilm. *Environ Sci Technol* **45**, 3367–3373 (2011). https://doi.org/10.1021/es103450g

14 Gulot, E. *et al.* Heterogeneity of diffusion inside microbial biofilms determined by fluorescence correlation spectroscopy under two-photon excitation. *Photochem Photobiol* **75**, 570–578 (2002). https://doi.org/10.1562/0031-8655(2002)075<0570:hodimb>2.0.co;2

15 Miller, J. K. *et al.* Nanoparticle deposition onto biofilms. *Ann Biomed Eng* **41**, 53–67 (2013). https://doi.org/10.1007/s10439-012-0626-0

16 Morrow, J. B., Arango, C. P. & Holbrook, R. D. Association of quantum dot nanoparticles with Pseudomonas aeruginosa biofilm. *J Environ Qual* **39**, 1934–1941 (2010). https://doi.org/10.2134/jeq2009.0455

17 Nevius, B. A., Chen, Y. P., Ferry, J. L. & Decho, A. W. Surface-functionalization effects on uptake of fluorescent polystyrene nanoparticles by model biofilms. *Ecotoxicology* **21**, 2205–2213 (2012). https://doi.org/10.1007/s10646-012-0975-3

18 Tripathi, S., Champagne, D. & Tufenkji, N. Transport behavior of selected nanoparticles with different surface coatings in granular porous media coated with Pseudomonas aeruginosa biofilm. *Environ Sci Technol* **46**, 6942–6949 (2012). https://doi.org/10.1021/es202833k

19 Jing, H. *et al.* Experimental and modeling studies of sorption of ceria nanoparticle on microbial biofilms. *Bioresour Technol* **161**, 109–117 (2014). https://doi.org/10.1016/j.biortech.2014.03.015

20 Li, X. *et al.* Control of nanoparticle penetration into biofilms through surface design. *Chem Commun (Camb)* **51**, 282–285 (2015). https://doi.org/10.1039/c4cc07737g

21 Xiao, Y. & Wiesner, M. R. Transport and retention of selected engineered nanoparticles by porous media in the presence of a biofilm. *Environ Sci Technol* **47**, 2246–2253 (2013). https://doi.org/10.1021/es304501n

22 Desmau, M. *et al.* Dynamics of silver nanoparticles at the solution/biofilm/mineral interface. *Environmental Science: Nano* **5**, 2394–2405 (2018). https://doi.org/10.1039/c8en00331a

23 Zhang, Z., Nadezhina, E. & Wilkinson, K. J. Quantifying diffusion in a biofilm of Streptococcus mutans. *Antimicrob Agents Chemother* **55**, 1075–1081 (2011). https://doi.org/10.1128/AAC.01329-10

24 Habimana, O. *et al.* Diffusion of nanoparticles in biofilms is altered by bacterial cell wall hydrophobicity. *Appl Environ Microbiol* **77**, 367–368 (2011). https://doi.org/10.1128/AEM.02163-10

25 Hollmann, B., Perkins, M., Chauhan, V. M., Aylott, J. W. & Hardie, K. R. Fluorescent nanosensors reveal dynamic pH gradients during biofilm formation. *npj Biofilms and Microbiomes* **7**, 50 (2021). https://doi.org/10.1038/s41522-021-00221-8

26 Rodríguez-Suárez, J. M., Gershenson, A., Onuh, T. U. & Butler, C. S. The heterogeneous diffusion of polystyrene nanoparticles and the effect on the expression of quorum-sensing genes and EPS production as a function of particle charge and biofilm age. *Environmental Science: Nano* **10**, 2551–2565 (2023). https://doi.org/10.1039/D3EN00219E
